# Functional Roles of Sensorimotor Alpha and Beta Oscillations in Overt Speech Production

**DOI:** 10.1101/2024.09.04.611312

**Authors:** Lydia Z. Huang, Yang Cao, Esther Janse, Vitória Piai

**Author notes:** Corresponding author: Lydia Z. Huang. Email address: Yang Cao, Esther Janse Vitória Piai.

## Abstract

Power decreases, or desynchronization, of sensorimotor alpha and beta oscillations (i.e., alpha and beta ERD) have long been considered as indices of sensorimotor control in overt speech production. However, their specific functional roles are not well understood. Hence, we first conducted a systematic review to investigate how these two oscillations are modulated by speech motor tasks in typically fluent speakers (TFS) and in persons who stutter (PWS). Eleven EEG/MEG papers with source localization were included in our systematic review. The results revealed consistent alpha and beta ERD in the sensorimotor cortex of TFS and PWS. Furthermore, the results suggested that sensorimotor alpha and beta ERD may be functionally dissociable, with alpha related to (somato-)sensory feedback processing during articulation and beta related to motor processes throughout planning and articulation. To (partly) test this hypothesis of a potential functional dissociation between alpha and beta ERD, we then analyzed existing intracranial electroencephalography (iEEG) data from the primary somatosensory cortex (S1) of picture naming. We found moderate evidence for alpha, but not beta, ERD’s sensitivity to speech movements in S1, lending supporting evidence for the functional dissociation hypothesis identified by the systematic review.

## 1 General Introduction

Speech motor processes are fundamental for daily spoken communication, and its dysfunction plays a pivotal role in neuromotor speech disorders. When one prepared the articulation of a word, the brain needs to plan and program the movements for the intended speech (Guenther et al., 1998). During articulation, the brain monitors the speech movements and exerts top-down control if they deviate from the intended trajectory (Tourville & Guenther, 2011). What are the neural markers associated with these processes and how do these neural markers differ in the case of neuromotor speech disorders? In this article, we intended to answer these questions by systematically reviewing neural oscillatory studies involving speech motor tasks in typically fluent speakers (TFS) and in persons who stutter (PWS). Furthermore, we followed up the systematic review findings with an intracranial electroencephalogram (iEEG) study to investigate whether the neural markers found in the review can be functionally dissociated.

Alpha and beta event-related desynchronization (ERD) are a prominent phenomenon in the sensorimotor areas during speech production. For speech movements, decreased power relative to the resting baseline is typically observed around sensorimotor cortex in alpha (8-13 Hz) and beta (15-25 Hz) frequency ranges before and throughout the movements (e.g., Jenson et al., 2014). Upon movement termination, power returns to baseline levels (e.g., Bowers et al., 2019; Gehrig et al., 2012), with beta oscillations around the precentral gyrus, specifically, showing strong power increase around 50 ms (De Nil et al., 2021), known as beta rebound or event-related synchronization (ERS).

Despite the increasing number of studies focusing on neural oscillations, the oscillatory markers of speech motor process have remained elusive. Speech production models have been primarily focused on the spatial location of processes (Hickok, 2012, 2014; Tourville & Guenther, 2011) drawing upon fMRI findings. Therefore, it is unclear which and how neural oscillations reflect speech motor processes before and during speech articulation. With the increasing number of neural oscillatory studies investigating this question (e.g., Bowers et al., 2019; Kittilstved et al., 2018), a systematic review of this literature is needed. The aims of the present article were to systematically review the evidence on alpha and beta oscillations for speech production in both speakers with fluent speech and speakers who stutter and to test empirically the patterns found in the review capitalizing on the high spatiotemporal resolution of iEEG data.

### 1 Systematic Review

For this systematic review, we were specifically interested in the associations between alpha and beta oscillations and speech motor function in fluent speakers and in speakers with stuttering. By reviewing studies that have used various electrophysiological techniques, we aimed to answer the following two research questions:

1. how are alpha and beta oscillations in canonical motor areas modulated by speech motor tasks?
2. do modulations of alpha and beta oscillations in speech motor tasks differ in PWS compared to TFS?

#### 1.1 Methods

Web of Science and PubMed databases were used to collect peer-reviewed studies between 1945 (1946 for PubMed) to 2021. These two databases were chosen for their good coverage of scientific journals, high paper quality, and relevance to medical and neuroscientific research. The database search was divided into two parts corresponding to our two research questions. For the first research question, the search was conducted using the keywords *motor* and *oscillation* and *speech production* (see Table S1 of the supplementary material). To include studies using different speech production paradigms, we further conducted a detailed search by crossing the terms *motor* and *oscillation* with one of the seven other terms that are subcategories of *speech production*: (1) *language production*, (2) *vowel production*, (3) *consonant production*, (4) *syllable production*, (5) *word production*, (6) *sentence production*, and (7) *articulation* (see Table S2 of the supplementary material). For the search terms with *articulation* in PubMed, we changed *oscillation* to *neural oscillation* to exclude studies on the oscillations of articulators. For the second research question, the terms *oscillation* and *stuttering* were used (see Table S3 of the supplementary material). Via these searches, 185 papers were identified after the removal of duplicates. To identify papers that were not detected in the automatic search, the reference lists from the included studies (see below) were screened. This did not yield additional papers. One additional study (De Nil et al., 2021) was included that was not identified via the above procedures.

##### Inclusion criteria

The study selection followed the following inclusion criteria (note that the criteria for studies on stuttering only differ in criterion 5 regarding participants):

1. The research topic of the paper should involve speech motor function;
2. The electrophysiological technique used in the study should be electro-encephalography (EEG), magneto-encephalography (MEG), electro-corticography (ECoG), and/or stereo-electroencephalography (SEEG);
3. The tasks in the experiments should involve at least one overt speech production task in which speech is elicited by having participants read or repeat words or phrases. Conceptually driven tasks, such as picture naming, were not considered because they involve long-term memory retrieval processes. Furthermore, for studies that compare between tasks, the tasks being compared should both involve speech processes. For example, if a study compared an overt speech production task with a non-speech tasks such as tongue protrusion, this study would not be considered in our review;
4. The time window of interest should include speech planning, articulation, or shortly following speech offset;
5. For the papers for the first research question, the recruited participants should be neurologically healthy TFS, whereas for the second research question, the participants should involve a group of PWS and a group of TFS;
6. The regions of interest (ROIs) should be at least one of the motor-related regions, including the sensorimotor cortex (SMC), the primary motor cortex (M1), the premotor cortex (PMC), and the supplementary motor area (SMA). Note that the primary somatosensory cortex (S1) was also considered in EEG and MEG studies considering its proximity to S1 and the low spatial resolution of surface measures. Furthermore, EEG and MEG studies should contain source-level analysis to ensure that the findings were optimally restricted to the ROIs;
7. The analysis should include frequency analysis of the oscillatory signals in alpha and/or beta bands.

Papers were screened by the first author (title and abstract in phase one and full-text in phase two) on whether they satisfied the above inclusion criteria. During the full-text screening, details on participants, ROIs, tasks, and analyzed time window were inspected and studies not meeting our inclusion criteria were removed, yielding 11 final studies (see Figure 1).

**Figure 1.**
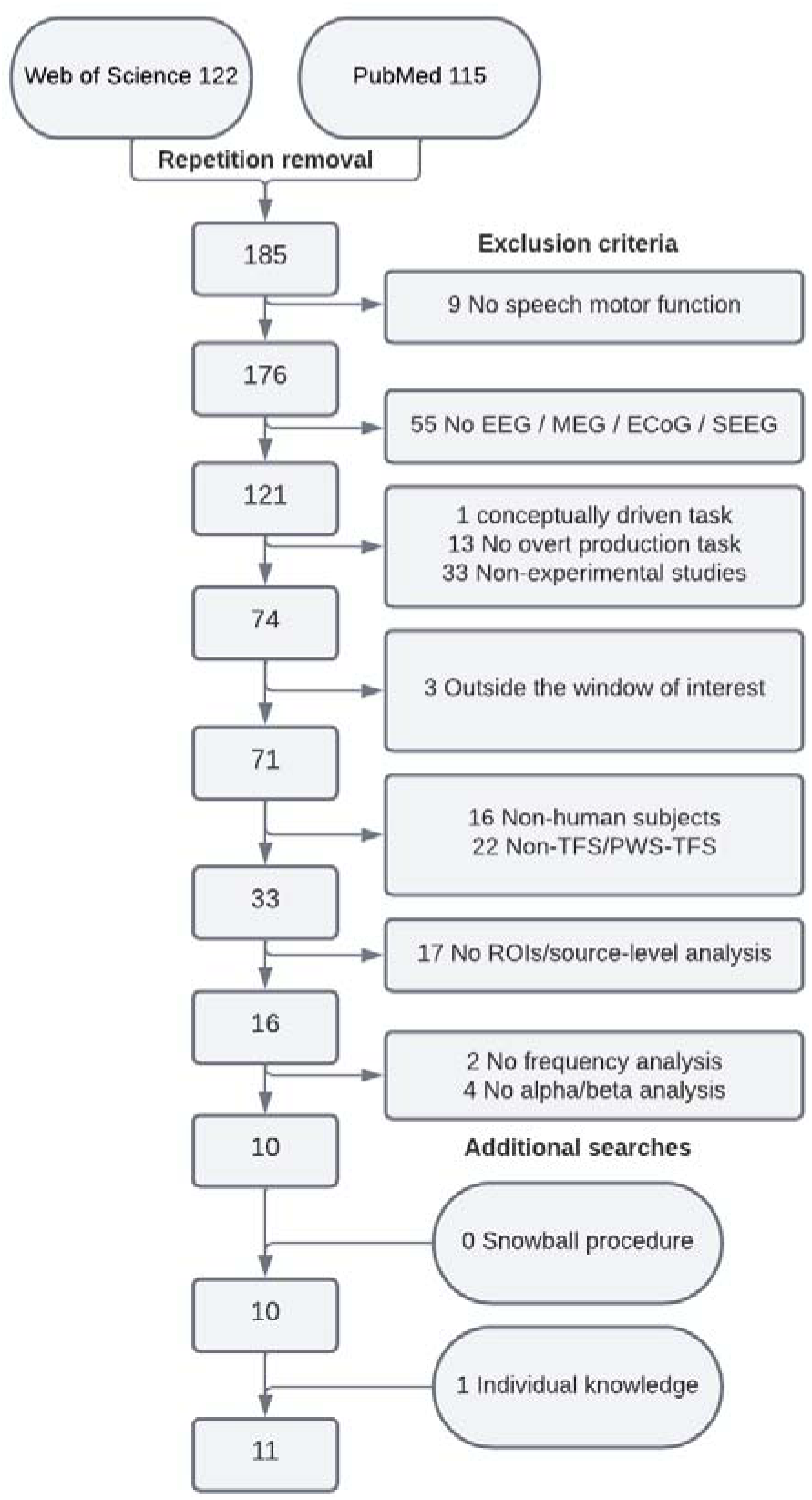
Literature Selection Process.

#### 1.2 Results

Descriptions of the 11 papers are shown in Tables 1 and 2. The six papers in Table 1 addressed the first research question, describing TFS’ sensorimotor alpha and beta patterns; the five papers in Table 2 focused on the second research question, which compared the sensorimotor alpha and beta patterns between PWS and TFS. All of these studies adopted surface measures (EEG/MEG) combined with source localization.

**Table 1.** Information of TFS studies (N = 6)

| Author<br>s | Year | Number of<br>subjects<br>(F/M) | Language | Study<br>Design | Conditions<br>(control<br>vs.<br>experimental) | Imaging<br>System | Multiple<br>Comparisons<br>Correction | Localization<br>Methods |
| --- | --- | --- | --- | --- | --- | --- | --- | --- |
| **EEG** |  |  |  |  |  |  |  |  |
| Bowers<br>et al.* | 2019 | 23<br>(20/3) | English | Delayed<br>syllable<br>reading | Cp vs. Op | 64-channel<br>NeuroScan<br>Quik<br>Cap | FDR | ICA, ECD,<br>BEM,<br>DIPFIT,<br>MARA |
| Jenson<br>et al. | 2014 | 20<br>(17/3) | English | Delayed<br>syllable<br>and<br>word<br>reading | Cp vs. Op | 68-channel<br>NeuroScan<br>Quik<br>Cap | FDR | ICA, ECD,<br>CSD,<br>sLORETA,<br>Talairach<br>cortical<br>atlas<br>digitized at<br>MNI,<br>BESA head<br>models |
| Kittilstved<br>et al. | 2018 | 26<br>(8/18) | English | Auditory<br>sentence<br>repetition | Op vs.<br>Choral /<br>DAF /<br>prolonged /<br>pseudo-stuttering | 68-channel<br>NeuroScan<br>Quik<br>Cap | FDR | ICA, ECD,<br>DIPFIT2,<br>Talairach<br>cortical<br>atlas,<br>BESA head<br>models,<br>MNI152<br>template |
| **MEG** |  |  |  |  |  |  |  |  |
| De Nil et al. | 2021 | 17 (9/8) | English | Delayed syllable reading | Simple Op vs. Complex Op | 151-channel CTF MISL | Bonferroni correction | MNI brain template, SPM8, event-related and frequency-based beamformer, a single sphere head model, Talairach coordinates, CIVET pipeline |
| Gehrig et al.* | 2012 | 26 (10/16) | German | Delayed sentence reading | Cp vs. Op | 275-channel CTF/VS M Omega | FWE | SPM8's 3D source reconstruction, LCMV beamformer |
| Gunji et al. | 2007 | 11 (4/7) | English | Delayed sentence reading | Singing vs. Op / humming / imagining singing | 151-channel CTF Omega | N/A, SAM permutation test | SAM beamformer, SPM T1 template, SPM99 |

**Table 2.** Information of PWS-TFS studies (N = 5)

| **Authors** | **Year** | **Number of subjects (F/M)** | **Language** | **Study Design** | **Conditions (control vs. experimental)** | **Imaging System** | **Multiple Comparisons Correction** | **Localization Method** |
| --- | --- | --- | --- | --- | --- | --- | --- | --- |
| **EEG** |  |  |  |  |  |  |  |  |
| Jenson et al. | 2020 | 11 stutter<br>11 control<br>(3/8;<br>3/8) | English | Auditory delayed syllable pair repetition | TFS vs. PWS | 128-channel Electrical Geodesics | Cluster-based corrections | ICA, principal component clustering, DIPFIT, MARA |
| Jenson et al. | 2018 | 31 stutter<br>24 control<br>(9/22;<br>7/17) | English | Delayed syllable pair and word reading | TFS vs. PWS | 68-channel NeuroScan QuikCap | Cluster-based corrections | ECD, DIPFIT2, PCA, sLORETA, Van Essen cortical maps, CSD, Talairach cortical atlas digitized at MNI, BESA head models, MNI152 brain template |
| **MEG** |  |  |  |  |  |  |  |  |
| Mersov et al. | 2016 | 12 stutter<br>12 control<br>(2/10;<br>2/10) | English | Delayed sentence reading | TFS vs. PWS | 151-channel CTF Omega | Tukey-HSD correction | SAM algorithm, BrainWave toolbox, MNI template, SPM8, MNI-Talairach conversion, |
|  |  |  |  |  |  |  |  | Talairach cortical atlas, CIVET pipeline |
| Mollaei et al. | 2021 | 11 stutter 11 control (2/9; 2/9) | English | Delayed sentence reading | TFS vs. PWS | 151-channel CTF Omega | TFCE | SAM algorithm, BrainWave toolbox, MNI standard space, AFNI, Monte Carlo simulations |
| Salmelin et al. | 2000 | 9 stutter 10 control (2/7; 2/8) | German | Delayed word reading | TFS vs. PWS | 122-channel Neuromag-122<sup>TM</sup> | N/A | TSE approach, a multi-dipole model, TSE curves comparison |

Note. PWS = people who stutter, TFS = typically fluent speakers, Cp = covert production, Op = overt production, Choral = Op with auditory accompaniment, DAF = Op with delayed auditory feedback, prolonged = prolonging the vowels in Op, pseudo-stuttering = pretending to stutter in Op. HSD = honestly significant difference, TFCE = threshold-free cluster enhancement, N/A = not applicable. AFNI = Analysis of neuroimages, from http://afni.nimh.nih.gov/afni/suma; BEM = Boundary element model; BESA = Cross-registration between spherical and realistic head geometry; CIVET = An image processing pipeline by Kim et al. (2015); CSD = Current source density; DIPFIT = A plugin at EEGLAB for localizing dipoles; ECD = Equivalent current dipole; ICA = Independent component analysis; LCMV = Linearly constrained minimum variance; MARA = Multiple artifact rejection algorithm; PCA = Principle component analysis; SAM = Synthetic aperture magnetometry; sLORETA = Standardized low-resolution brain electromagnetic tomography; SPM = Statistical parametric mapping
\* Papers with connectivity measures.

##### 1.2.1 Sensorimotor Alpha and Beta Patterns in Typically Fluent Speakers

In summary, most participants in the TFS studies were adults with a few teenagers at the age of 16. The presented stimuli in all TFS studies were either a sentence, a word, or a syllable pair. Note that the stimuli were presented visually except in Kittilstved et al. (2018), where auditory stimuli were used and participants were instructed to repeat them aloud. To extract the motor component in the task, most TFS studies contrasted overt production (OP) with covert production (CP) with CP as a control condition (Bowers et al., 2019; Gehrig et al., 2012; Gunji et al., 2007; Jenson et al., 2014; see Table 1). Participants were required to read the (visually) presented stimulus aloud in the OP condition and to imagine speaking the stimulus aloud in the CP condition. Other TFS studies used OP as a control condition and compared it with other OP conditions in which task instructions or sensory conditions were manipulated, such as requiring the participants to pretend to stutter or delaying the auditory feedback during OP (De Nil et al., 2021; Gunji et al., 2007; Kittilstved et al., 2018). Alpha and beta ERDs were consistently found during OP in motor-related cortices through statistical testing or visual inspection. The observed areas included the PMC, M1, S1, SMA, and SMC. Details of these alpha and beta ERD sources per study are shown in Table 1. These sensorimotor alpha and beta ERDs occurred following stimulus/cue onset (Bowers et al., 2019; De Nil et al., 2021; Gehrig et al., 2012; Jenson et al., 2014; Kittilstved et al., 2018), and were sustained throughout articulation (Bowers et al., 2019; De Nil et al., 2021; Gunji et al., 2007; Kittilstved et al., 2018). Additionally, a beta rebound (i.e., increased power) was found around 50 ms following articulation offset (De Nil et al., 2021). Below, we describe the results from individual studies in more detail.

Three studies compared alpha and beta power in OP relative to CP in motor-related cortices. With EEG, Bowers et al. (2019) observed significantly greater beta ERD in the SMC bilaterally throughout planning and articulation for syllable production, while greater alpha ERD was observed only in the planning stage in the bilateral SMC. Jenson et al. (2014) also found greater beta ERD, but not alpha ERD, in the planning stage in the bilateral SMC and PMC for syllable and word production. With MEG, Gehrig et al. (2012) found stronger beta but no alpha ERD in the left M1 and right S1 during planning for sentence reading; stronger alpha ERD was found in the SMA. These results associate the sensorimotor alpha ERD with processes primarily in the articulation stage and the sensorimotor beta ERD with processes that span across speech planning and articulation.

The other three studies (De Nil et al., 2021; Gunji et al., 2007; Kittilstved et al., 2018) included additional sensory manipulations or specific speech materials or instructions in the OP task and compared these adjusted tasks with the normal OP condition (simply speaking aloud). De Nil et al. (2021) manipulated speech materials, and specifically investigated whether beta power is sensitive to articulatory complexity throughout syllable production. They found a beta power difference during planning and after articulation of complex relative to simple syllables (e.g., /pa ta ka pa/ vs. /pa pa pa pa/). Stronger beta ERD was observed in the bilateral SMC before articulation, and stronger beta ERS was found only in the left SMC after articulation offset. These findings suggest that the sensorimotor beta ERD is sensitive to increasing motor demands.

Kittilstved et al. (2018) manipulated speech instructions, as well as timing of auditory feedback, and examined the sensitivity of alpha and beta power in the SMC to modulated motor or sensory processing effort in sentence repetition. One task required additional motor demands by instructing participants to pretend to stutter during articulation, namely *pseudo-stuttering* (authors’ own terminology). Two additional tasks with modulated sensory demands were included, which respectively required participants to articulate a sentence when receiving delayed auditory feedback or to prolong the vowels in the sentence (conditions “DAF” and “prolonged”, authors’ own terminology). While DAF invokes monitoring of delayed external sensory signal, the prolonged condition demands the participants to monitor the increased feedback during articulation. The time window upon speech onset was inspected. Stronger beta ERD was observed in the left SMC for pseudo-stuttering relative to OP. Greater alpha ERD was observed in the right SMC for both the DAF and prolonged conditions relative to OP. These results indicate that the sensorimotor beta ERD is sensitive to increased motor demands and the sensorimotor alpha ERD to increased sensory demands. Note that Kittilstved et al. (2018) also included a choral speech condition in their study, in addition to the conditions described above. However, we excluded this condition for its lack of comparability in design to the control condition of normal repetition, as participants were exposed to the stimulus visually earlier in the choral than control condition. By contrast, the experimental conditions we discussed above were comparable to the control condition without any obvious confounds.

Gunji et al. (2007) provided supporting evidence for sensorimotor alpha’s sensitivity to sensory processing by including the instruction to sing. They found significantly stronger alpha ERD in the right SMC in singing compared to speaking and humming the same sentence, “Happy birthday to you”. This finding was interpreted to reflect the additional demand for controlling the melody during articulation, namely, to process and maintain the pitch of tone in singing compared to speaking and humming. However, these results should be considered more critically since the data were corrected by a post-articulation baseline, a 7-s time window immediately following articulation offset. This time window is known to contain residual brain activity related to speech movement, including beta rebound and alpha power returning to baseline (e.g., Bowers et al., 2019; De Nil et al., 2021).

##### 1.2.2 Sensorimotor Alpha and Beta Deviance in Stuttering

In summary, the participants in the PWS-TFS studies were predominantly age- and sex-matched adult PWS and TFS speakers, ranging from 17 to 55 years old. The stimuli in the PWS-TFS studies were either a sentence, a word, or a syllable pair. Most studies presented all of their stimuli visually (Jenson et al., 2018; Mersov et al., 2016; Salmelin et al., 2000), except Jenson et al. (2020), which presented their stimuli auditorily. The most common paradigm was OP in which the PWS group was directly contrasted with the TFS group (Jenson et al., 2018; Mersov et al., 2016; Salmelin et al., 2000). Participants were instructed to read the (visually) presented stimulus aloud. Only one study, Jenson et al. (2018), used CP as a control condition for OP before comparing the PWS with TFS groups. Alpha and beta ERDs were found through statistical testing or visual inspection in sensorimotor areas, including PMC, M1, and S1, across TFS and PWS during planning and articulation of OP (for details of these sources, see Table 2). However, whether groups differed, and if so, how, varied across studies. These mixed findings may relate to between-study differences in electrophysiological techniques, stuttering severity, and speech stimuli. More details on these mixed findings of the sensorimotor alpha and beta ERDs are provided below.

###### Alpha ERD

Two MEG studies failed to observe significant between-group differences in alpha power for sentence and word production, respectively (Mersov et al., 2016; Salmelin et al., 2000). Two EEG studies, by contrast, found reduced alpha ERD in PWS vs. TFS for syllable production (including CP and OP) and overt word production (Jenson et al., 2018, 2020).

However, the findings in overt syllable production were inconsistent between the two EEG studies. While Jenson et al. (2020) observed a significantly reduced alpha ERD bilaterally in syllable production, Jenson et al. (2018) only observed this phenomenon in word but not in syllable articulation in the left hemisphere of PWS vs. TFS. One possible cause for this difference is the varying stuttering severity of PWS included in these two studies (Jenson et al., 2020: moderate to severe vs; Jenson et al., 2018: mostly mild). The prominent reduced alpha ERD in Jenson et al. (2020) may be related to the relatively more severe stuttering severity of their PWS group. Another possible explanation for the different findings is the different stimulus modality (auditory vs. visual presentation). However, it is unlikely that early perceptual differences would influence the sensorimotor alpha ERD, which is observed at a much later cognitive stage.

###### Beta ERD

Mixed findings were found in beta ERD when comparing PWS with TFS. Overall, one EEG study reported reduced beta ERD for syllable production (including covert and overt) and overt word production (Jenson et al., 2018; Jenson et al., 2020 did not find significant effects, but their sample was smaller), while two MEG studies reported increased beta ERD relative to baseline either in the mouth (Mersov et al., 2016) or the hand motor area (Salmelin et al., 2000). These findings were primarily localized to the PMC and M1 (see Table 2). Salmelin et al. (2000) did not find significant between-group differences in beta ERD in the mouth motor area; however, they found significantly stronger beta ERD in the hand motor area for PWS relative to TFS for word production. They suggest that it may reflect less clear functional dissociation of hand and mouth areas in PWS compared to TFS, but the authors could not rule out the possibility that this effect was induced by excessive hand movements of PWS. They found a trend of reduced beta ERD in PWS compared to TFS in the results of the mouth motor area, consistent with the pattern observed in the EEG studies. By contrast, Mersov et al. (2016) found bilaterally stronger beta ERD in PWS relative to TFS, opposite to the findings of Salmelin et al. (2000). Only the stronger beta ERD in the left BA6 survived correction for multiple comparisons (Mersov et al., 2016).

These observed discrepancies in the modulation of beta ERD may be related to differences in stimulus type across studies. While the studies that observed reduced ERD (Jenson et al., 2018, 2020; Salmelin et al., 2000) employed relatively short stimuli, such as words or syllable pairs, participants in Mersov et al. (2016) were required to read a sentence aloud.

#### 1.3 Discussion

In sum, relating to our first research question, we found consistent sensorimotor alpha and beta ERDs during planning and articulation for overt speech production with varying lengths of stimuli and ERSs after articulation offset. The sensorimotor alpha ERD presents sensitivity to the articulation stage for OP relative to CP bilaterally and to sensory-feedback demanding tasks relative to ‘normal’ OP in the right hemisphere. By contrast, the sensorimotor beta ERD is sensitive to both planning and articulation for OP relative to CP and to motor-demanding tasks primarily in the left hemisphere. These findings from healthy fluent speakers were consistently observed among TFS and PWS. However, relating to our second research question, the results from contrasting PWS and TFS were less consistent. Weaker sensorimotor alpha ERD for PWS relative to TFS was observed during articulation in the two EEG studies but absent in the two MEG studies. Furthermore, while weaker beta ERD for PWS was observed primarily in the EEG studies, stronger beta ERD was found in one MEG study. These findings were observed throughout planning and articulation of OP.

Some of the observed discrepancies in stuttering studies could be the result of EEG-MEG differences. MEG is mostly sensitive to tangential sources while EEG to both tangential and radial sources (Hari et al., 2010; Singh, 2014). The MEG signal is also less affected by motor artifacts compared to EEG due to relatively less volume conduction. In addition, different localization methods and data processing approaches were used across the studies included in our systematic review, which could muddle the picture further.

Stimulus type could also be a potential cause for some of the observed discrepancies in both the PWS and TFS studies. It is noticeable that studies with sentence reading paradigms (Gehrig et al., 2012; Mersov et al., 2016) are generally inconsistent with other comparable studies with a word/syllable reading paradigm. To recap, while Gehrig et al. (2012) observed stronger beta ERD in the left M1 and right S1 for planning of OP relative to CP, other studies focusing on the same OP-CP contrast found the beta ERD pattern bilaterally. Similarly, Mersov et al. (2016) found stronger beta ERD in the left PMC for PWS relative to TFS; other similar studies observed reduced beta ERD in the PMC and M1 bilaterally. Although the complex pathology of stuttering may add to the discrepancies among the studies on stuttering, the specific modulation in the left hemisphere corresponds well with the typical findings in sentence production being highly left-lateralized (Lukic et al., 2021; Tremblay & Small, 2011).

The variability in stimulus type may not only lead to mixed results in the review findings but may also limit the generalizability of our results from TFS studies. For example, reviewed studies that manipulated motoric and sensory factors in OP primarily adopted a sentence reading paradigm (Gunji et al., 2007; Kittilstved et al., 2018), which produced highly lateralized results. Specifically, the sensory-demanding task modulated the alpha ERD in the right SMC, while a motor-demanding task affected the beta ERD in the left SMC. By contrast, De Nil et al. (2021) found bilateral modulation of beta ERD in syllable production of syllables varying in complexity, and hence, varying motor demands. Replication of these studies with varying stimulus types is needed to generate a clear picture of these task effects.

To the best of our knowledge, this is the first review to systematically examine how sensorimotor alpha and beta oscillations are modulated by speech motor tasks. Previous reviews on similar topics were neither systematic nor focused on the sensorimotor alpha and beta oscillations in overt speech production (e.g., Jenson et al., 2020; Piai & Zheng, 2019; Prystauka & Lewis, 2019). Although Jenson et al. (2020) zoomed in on the sensorimotor aspect of the mu rhythm, their findings were drawn from a wide range of cognitive studies that compared PWS with TFS. Our review suggests that future replication that takes the potential role of stuttering severity into account is needed to fully understand the roles of sensorimotor alpha and motor oscillations among PWS.

Our findings reveal a tentative pattern of a functional dissociation between the sensorimotor alpha and beta ERDs in speech planning and articulation. Tentatively, the sensorimotor alpha ERD seems to reflect (somato-)sensory feedback processing as it showed sensitivity to changing sensory feedback. The finding of stronger alpha ERD in the articulation but not planning stage of OP relative to CP implicates the same functional association, following the logic that overt but not covert speech movements generate somatosensory feedback for articulation. By contrast, the sensorimotor beta ERD is consistently modulated by changing motor demands and is sensitive to the continuous and greater demands for motor planning, programming, and control throughout planning and articulation of OP relative to CP. These findings from the systematic review feed directly into hypotheses, which we tested in an iEEG study, to which we turn next.

## 2 iEEG Study

From the findings described above and considering the canonical roles of S1 and M1 within the SMC, we hypothesized a functional-anatomical dissociation of the sensorimotor alpha and beta ERDs related to speech production. Functionally, alpha ERD is likely related to somatosensory feedback processing while beta ERD is associated with motor processes. Following the functionally delineated roles of S1 and M1, alpha and beta ERDs are hypothesized to show distinct modulation in these two areas, that is, with a distinct pattern of alpha and beta ERDs in S1 and M1 during speech planning and articulation. The alpha ERD is expected to be stronger during the articulation than the planning stage in S1 but relatively unchanged in M1. This prediction corresponds to the demands for processing the somatosensory feedback only present during the articulation but not the planning stage and the canonical role of S1 but not M1 in this process. By contrast, the beta ERD is expected to remain at a similar level throughout the planning and articulation stages in M1 and S1. The beta ERD in M1 should be stronger than in S1 as motor processes are required in both the planning and articulation stages and canonically occur at M1 but not S1. Using existing iEEG data from S1 during overt word production, we tested the prediction that alpha ERD would be stronger during articulation than preceding it, whereas beta ERD would be similar across the two stages. Available data on M1 was limited, preventing us from testing hypotheses regarding M1.

Traditional frequency analysis (i.e., narrowband filtering) assumes specific neural oscillatory band boundaries and the presence of activity in ROIs while ignoring the presence of confounding non-oscillatory aperiodic activity (Donoghue et al., 2022). Studies have shown that aperiodic components and oscillatory frequency vary across participants and cognitive states (e.g., Donoghue et al., 2020; Grandy et al., 2013; Haegens et al., 2014). In our analysis, we took these methodological considerations into account. Specifically, we verified the existence of oscillations and validated the oscillatory band boundaries, and considered the concurrent non-oscillatory aperiodic activity to minimize potential confounds by using fitting oscillations and the one-over-f (FOOOF) algorithm (Donoghue et al., 2020). The FOOOF algorithm allows us to parameterize the periodic and aperiodic components in an individual power spectrum.

### 2.1 Method

Existing, available human iEEG data of spoken word production from Woolnough et al. (2022) were analyzed. Given anatomical coverage limitations in M1, we focused on alpha and beta oscillations in S1. Details of the experimental design, implantation and data collection can be found in Woolnough et al. (2022). The original data were downloaded from the Data Archive for the BRAIN Initiative (DABI) in Neurodata Without Borders Version 2.2.5 format.

#### 3.1.1. Participants

iEEG recordings from eight patients out of 50 patients (52 recordings) in Woolnough et al. (2022) were selected and analyzed. The selected patients had subdural or depth electrodes implanted in S1 for seizure localization of refractory epilepsy. Details of these patients and bipolar contacts are shown in Table 5.

**Table 3.**
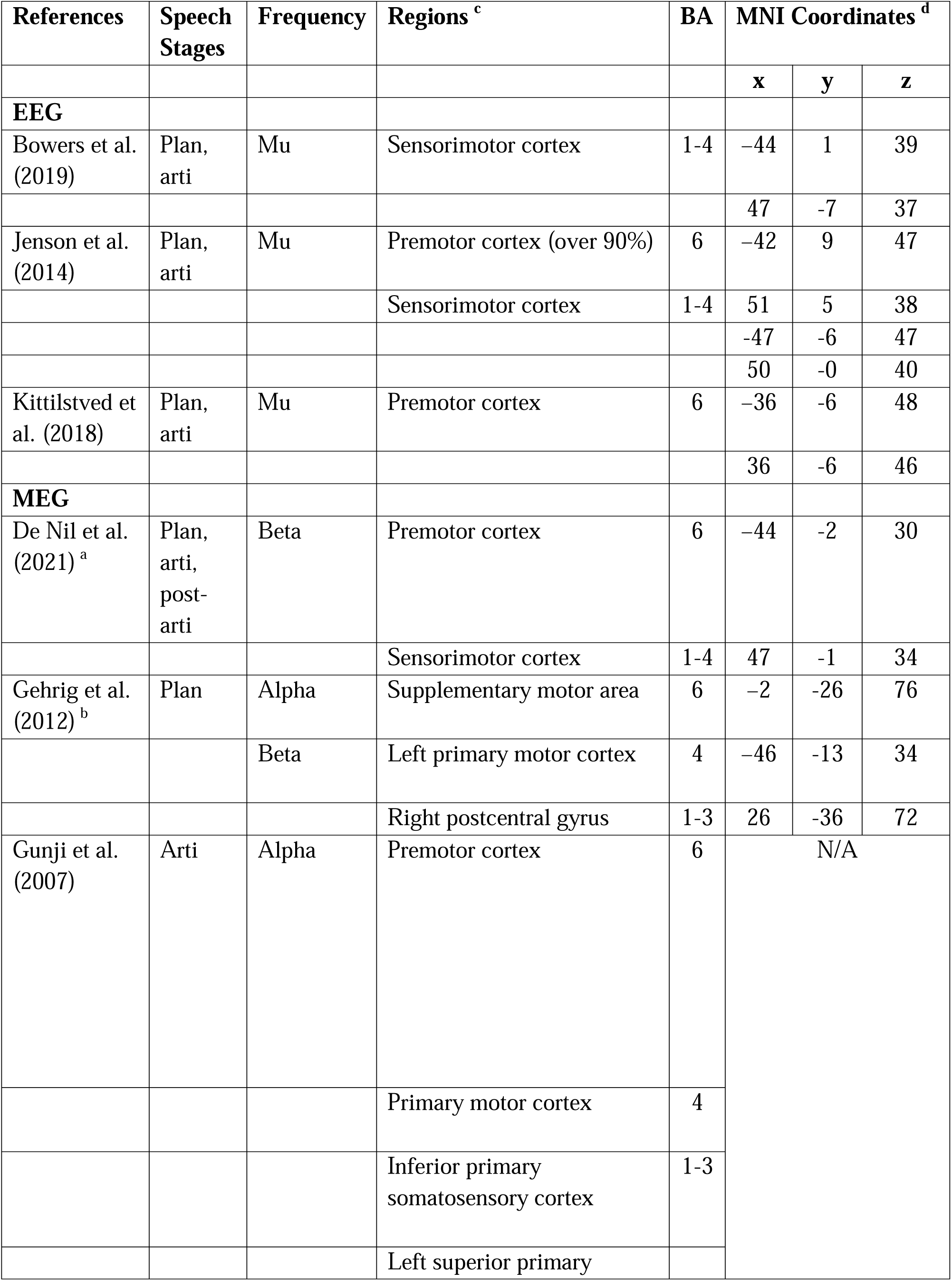
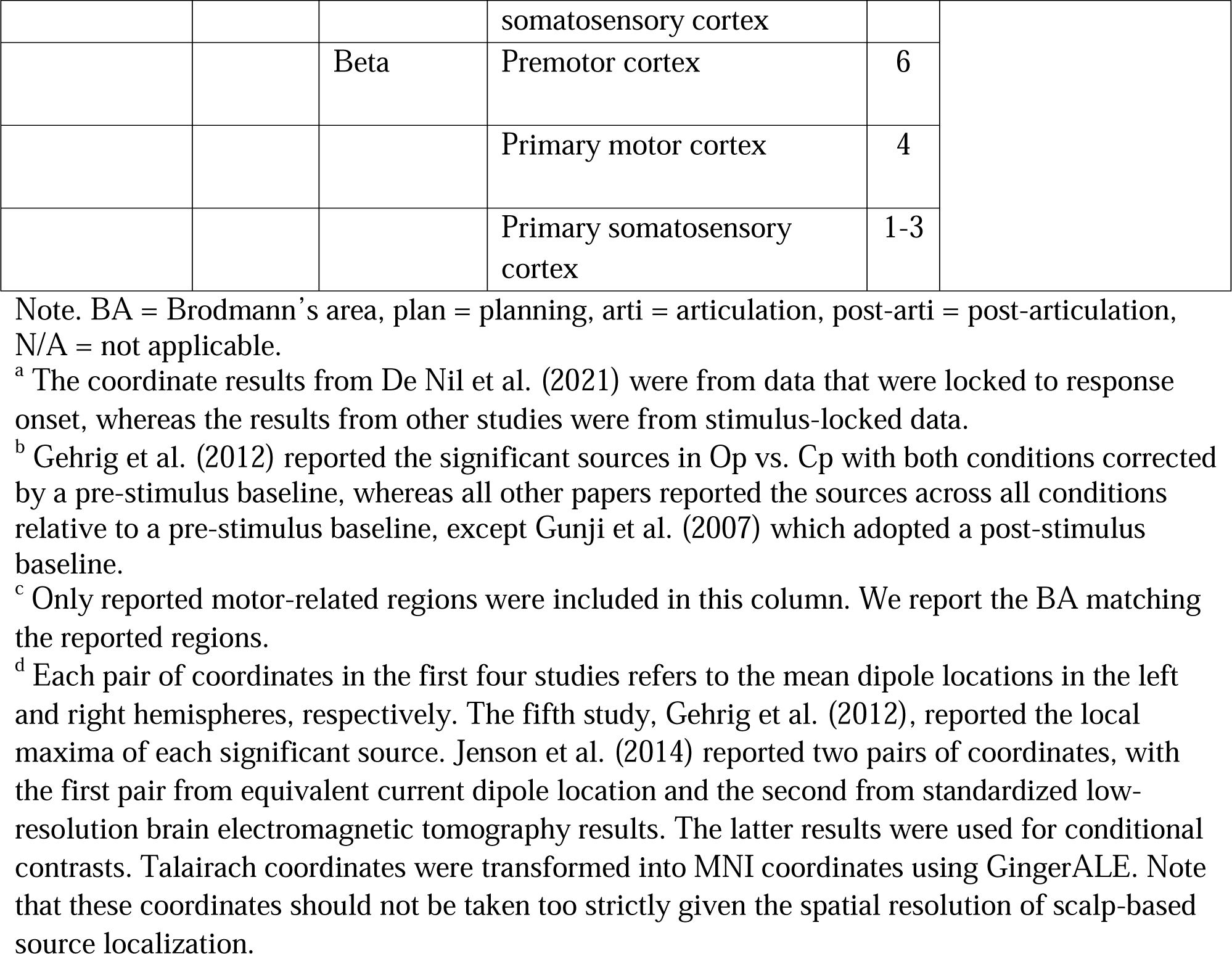
Localization of Sensorimotor-Related Alpha and Beta Power Changes in TFS Studies.

| References | Speech Stages | Frequency | Regions <sup>c</sup> | BA | MNI Coordinates <sup>d</sup> |  |  |
| --- | --- | --- | --- | --- | --- | --- | --- |
|  |  |  |  |  | x | y | z |
| EEG |  |  |  |  |  |  |  |
| Bowers et al. (2019) | Plan, arti | Mu | Sensorimotor cortex | 1-4 | −44 | 1 | 39 |
|  |  |  |  |  | 47 | -7 | 37 |
| Jenson et al. (2014) | Plan, arti | Mu | Premotor cortex (over 90%) | 6 | −42 | 9 | 47 |
|  |  |  | Sensorimotor cortex | 1-4 | 51 | 5 | 38 |
|  |  |  |  |  | -47 | -6 | 47 |
|  |  |  |  |  | 50 | -0 | 40 |
| Kittilstved et al. (2018) | Plan, arti | Mu | Premotor cortex | 6 | −36 | -6 | 48 |
|  |  |  |  |  | 36 | -6 | 46 |
| MEG |  |  |  |  |  |  |  |
| De Nil et al. (2021) <sup>a</sup> | Plan, arti, post-arti | Beta | Premotor cortex | 6 | −44 | -2 | 30 |
|  |  |  | Sensorimotor cortex | 1-4 | 47 | -1 | 34 |
| Gehrig et al. (2012) <sup>b</sup> | Plan | Alpha | Supplementary motor area | 6 | −2 | -26 | 76 |
|  |  | Beta | Left primary motor cortex | 4 | −46 | -13 | 34 |
|  |  |  | Right postcentral gyrus | 1-3 | 26 | -36 | 72 |
| Gunji et al. (2007) | Arti | Alpha | Premotor cortex | 6 | N/A |  |  |
|  |  |  | Primary motor cortex | 4 |  |  |  |
|  |  |  | Inferior primary somatosensory cortex | 1-3 |  |  |  |
|  |  |  | Left superior primary |  |  |  |  |

|  |  |  |  |  |
| --- | --- | --- | --- | --- |
|  |  |  | somatosensory cortex |  |
|  |  | Beta | Premotor cortex | 6 |
|  |  |  | Primary motor cortex | 4 |
|  |  |  | Primary somatosensory cortex | 1-3 |

Note. BA = Brodmann's area, plan = planning, arti = articulation, post-arti = post-articulation, N/A = not applicable.
<sup>a</sup> The coordinate results from De Nil et al. (2021) were from data that were locked to response onset, whereas the results from other studies were from stimulus-locked data.
<sup>b</sup> Gehrig et al. (2012) reported the significant sources in Op vs. Cp with both conditions corrected by a pre-stimulus baseline, whereas all other papers reported the sources across all conditions relative to a pre-stimulus baseline, except Gunji et al. (2007) which adopted a post-stimulus baseline.
<sup>c</sup> Only reported motor-related regions were included in this column. We report the BA matching the reported regions.
<sup>d</sup> Each pair of coordinates in the first four studies refers to the mean dipole locations in the left and right hemispheres, respectively. The fifth study, Gehrig et al. (2012), reported the local maxima of each significant source. Jenson et al. (2014) reported two pairs of coordinates, with the first pair from equivalent current dipole location and the second from standardized low-resolution brain electromagnetic tomography results. The latter results were used for conditional contrasts. Talairach coordinates were transformed into MNI coordinates using GingerALE. Note that these coordinates should not be taken too strictly given the spatial resolution of scalp-based source localization.

**Table 4.** Localization of Sensorimotor-Related Alpha and Beta Power Changes in PWS-TFS Studies.

| References | Frequency | Regions | BA | MNI Coordinates |
| --- | --- | --- | --- | --- |

|  | a |  |  | b |  |  |
| --- | --- | --- | --- | --- | --- | --- |
|  |  |  |  | x | y | z |
| **EEG** |  |  |  |  |  |  |
| Jenson et al. (2018) | Alpha ↓<br>Beta ↓ | Left premotor cortex (ECD) | 6 | -35 | -4 | 45 |
|  |  | Right premotor cortex (ECD) | 6 | 37 | -2 | 47 |
|  |  | Left primary motor cortex (sLORETA) | 4 | -41 | -11 | 53 |
|  |  | Right premotor cortex (sLORETA) | 6 | 39 | -5 | 52 |
| Jenson et al. (2020) | Alpha ↓ | Primary motor cortex | 6 | N/A |  |  |
|  |  | Primary somatosensory cortex | 1-3 |  |  |  |
| **MEG** |  |  |  |  |  |  |
| Mersov et al. (2016) | Beta ↑ | Ventral premotor cortex | 6 | -48 | 2 | 27 |
|  |  |  |  | 55 | 7 | 29 |
| Salmelin et al. (2000) | Beta ↑ | Primary motor cortex | N/A | N/A |  |  |

Note. BA = Brodmann's area, ECD = equivalent current dipole, sLORETA = standardized low-resolution brain electromagnetic tomography, N/A = not applicable. The localization results were estimated across all participant groups and conditions relative to a pre-stimulus baseline, except that Mersov et al. (2016) separately localized the sources of TFS and PWS groups; and both groups produced the same results.
<sup>a</sup> The upward- and downward-pointing arrows next to "alpha" and "beta" represent the direction of alpha and beta ERDs in PWS compared to TFS. Upward indicates stronger ERD in PWS relative to TFS, and vice versa.
<sup>b</sup> Each pair of coordinates in each study refers to the mean dipole locations in the left and right hemispheres, respectively. Talairach coordinates were transformed into MNI coordinates using GingerALE. Note that these coordinates should not be taken too strictly given the spatial resolution of scalp-based source localization.

**Table 5.** Patients’ Information.

| Patient | Age | Sex | Hand | Electrode | Hemisphere | Bipolar | Num |
| --- | --- | --- | --- | --- | --- | --- | --- |

|  |  |  |  | type |  | contacts | ber of trials available for analysis |
| --- | --- | --- | --- | --- | --- | --- | --- |
| TA525 | 42 | F | L | SDE | R | S1-S1 | 34 |
| TA530 | 34 | F | R | SDE | L | S1-a9-46v | 24 |
| TA533 | 29 | M | L | SDE | R | S1-S1 | 48 |
| TA641 | 45 | F | R | SEEG | L | S1-S1 | 62 |
| TA774 | 29 | F | L | SEEG | R | S1-S1 | 30 |
| TS024 | 31 | F | R | SDE | L | S1-S1 | 37 |
| TS060C | 37 | M | R | SDE | L | PFm-S1 | 39 |
| TS081 | 18 | F | R | SDE | R | S1-S1 | 48 |

Note. The locations of the contacts include primary somatosensory cortex (S1), inferior parietal cortex (PFm), dorsolateral prefrontal cortex (a9-46V). M = male, F = female, L = left, R = right, SDE = subdural electrode, SEEG = stereo-electroencephalography electrode.

#### 3.1.2. Experimental design

All subjects completed a landmark identification task in which they were presented with a color photograph of a famous landmark and were asked to recall the location of this landmark and name it aloud. Given that we made use of an existing dataset, the choice of task was determined by the original investigation. As a control condition in the original study (but not used in the present study), all subjects were also presented with a scrambled version of the famous landmark, and they were instructed to say “scrambled” aloud. Coherent images and their scrambled versions were presented in a pseudorandomized order. A total of 140-160 pictures were presented in each recording session. Each picture was displayed for 2000 ms followed by an inter-stimulus interval of 6000 ms. Offline, the accuracy and response onset of each trial were manually marked based on audio recordings. Responses that matched the city, state (province), or country of the landmark were considered correct. For the present study, we only analyzed the correct trials from the coherent condition for its resemblance to a standard language production task such as picture naming. This also avoids potentially confounding effects from the stereotypical production of “scrambled” (Bullock et al., 2023).

#### 3.1.3. Contact and data selection

Data analyzed in the present study were obtained from either subdural grid electrodes (SDEs; 6 patients) or stereotactically implanted depth electrodes (sEEGs; 2 patients). We devised criteria for selecting one pair of contacts in the region of interest per patient, as follows.

We first selected contacts with the Human Connectome Project label 3, 1, or 2, corresponding to S1. For patients with only one S1 contact (TA530 and TS060C), we additionally selected a contact adjacent to the S1 contact (see Table 5). Upon obtaining the set of usable S1 contacts, we excluded contacts that were outside of the ventral and dorsal S1 areas (i.e., z < 56 using coordinates provided with the shared dataset). This exclusion criterion was based on the meta-analysis of the term “articulatory” in Neurosynth (https://neurosynth.org/, Yarkoni et al., 2011). We further inspected contacts labelled as “good” in the dataset trial by trial to exclude those with no activity recorded. Finally, considering that our review findings generally involve the ventral SMC (Bowers et al., 2019; De Nil et al., 2021; Gehrig et al., 2012), we selected the two adjacent contacts that were the most ventral per patient. This procedure resulted in one pair of contacts per patient with the most optimal a-priori defined location.

We extracted stimulus-locked trials and their response-locked correspondents for the contacts, condition (i.e., landmark naming), and trials (i.e., correct responses) of interest. Finally, trials that required monosyllabic responses were excluded to avoid including trials with speech duration shorter than our time window of analysis (i.e., 400 ms, see below), as beta rebound could occur after articulation offset. This resulted in 40.25 available trials per patient on average.

#### 3.1.4. Data processing and statistical analysis

Our offline data processing was conducted using FieldTrip (Oostenveld et al., 2011) in Matlab R2021a (The MathWorks, Natick, MA) and FOOOF (version 1.0.0, Donoghue et al., 2020) in Python 3.8 (Python Software Foundation, https://www.python.org/).

First, per patient and contact, baseline segments were extracted from the trial-level data from −400 to 0 ms locked to stimulus onset, and response-locked segments were created from −1 to 2 s locked to speech onset, based on the naming RT. Second, the response-locked segments underwent baseline correction, in which the time-average of the baseline segment data was subtracted from each data time point.

Next, bipolar re-referencing and frequency analysis were performed. For each pair of contacts in each patient, the data points in one contact were subtracted from those in the other contact. Before frequency analysis, the response-locked segment was further divided into two segments: the *planning* segment, which was 400 ms before speech onset, and the *articulation* segment, which was 400 ms starting at speech onset. The baseline, planning and articulation segments (henceforth collectively termed “trial events“) underwent spectral decomposition with the fast Fourier transform (FFT) between 2 and 40 Hz in steps of 1 Hz using a Hanning taper.

This FFT analysis was performed once on the segments separately to obtain the power spectra for each trial event and once over all trial events combined. Note that the 400-ms window was chosen to balance between the frequency resolution and the RTs of responses with bi- or trisyllabic words observed in the data (> 400 ms) and their typical speech duration (> 400 ms, Jacewicz et al., 2009, 2010).

Per participant and bipolar contact, the FOOOF algorithm (version 1.0.0, Donoghue et al., 2020) was used to parameterize the power spectra obtained from the spectral decomposition step. Default settings of the algorithm were used except that the frequency range was set between 2 and 40 Hz. We first extracted and inspected the dominant peak frequencies and peak power from the parametrized power spectra for each trial event. Overall, three peaks were found in the power spectra for each trial event: an alpha peak at 8 Hz on average, a low beta peak at 19 Hz on average, and a high beta peak at 25 Hz on average. The first two peaks were more commonly found than the high beta peak among all trial events and eight participants (patient TA530’s planning and articulation events showed no parameterized alpha peak; the high beta peak was only parameterized in the baseline event of three participants and therefore not analyzed further). As a result, we extracted power at the relatively consistent alpha and low beta peak frequencies for each trial event. To determine the alpha power in TA530’s planning and articulation events with missing peak frequencies, we used the dominant alpha frequency across all trial events (i.e., 8 Hz) as a proxy. Finally, the extracted peak power of the planning and articulation events were baseline normalized, using the formula: 100* ((condition – baseline)/baseline)).

For the statistical analyses, Bayesian t-tests were conducted on alpha and beta bands separately, using JASP (version 0.17.1.0). First, to verify the existence of ERD, a one-sample t-test was conducted for each speech stage, with a test value of 0 and an alternative hypothesis that the power values are smaller than the test value (i.e., power is decreased relative to baseline). To test our a-priori hypotheses, a paired-samples t-test was used on the dataset from each frequency band. Within each test, the data were inspected collectively for outliers (IQR >1.5), resulting in the exclusion of one participant (TA774) for their extreme values in the alpha band, yielding a final sample size of seven patients. Then, the normality of the paired differences was checked by applying a Shapiro-wilk test to the difference in values between planning and articulation stages. The normality assumption was not violated. Two hypotheses were then respectively tested in each of the tests: 1) alpha ERD is stronger during articulation than during planning, 2) beta ERD is similar during articulation and planning. Descriptive statistics and their plots, prior and posterior, Bayes factor robustness check, and sequential analysis were inspected (Goss-Sampson et al., 2020).

### 2.2 Results

From the final seven included patients, the dominant alpha frequency was at 9 Hz and beta at 18 Hz, on average. Power changes relative to baseline in the speech planning and articulation stages for the alpha and beta bands for each individual patient is shown in Figure 2. Fitted power spectra at the individual level is shown in Figure S1.

**Figure 2.**
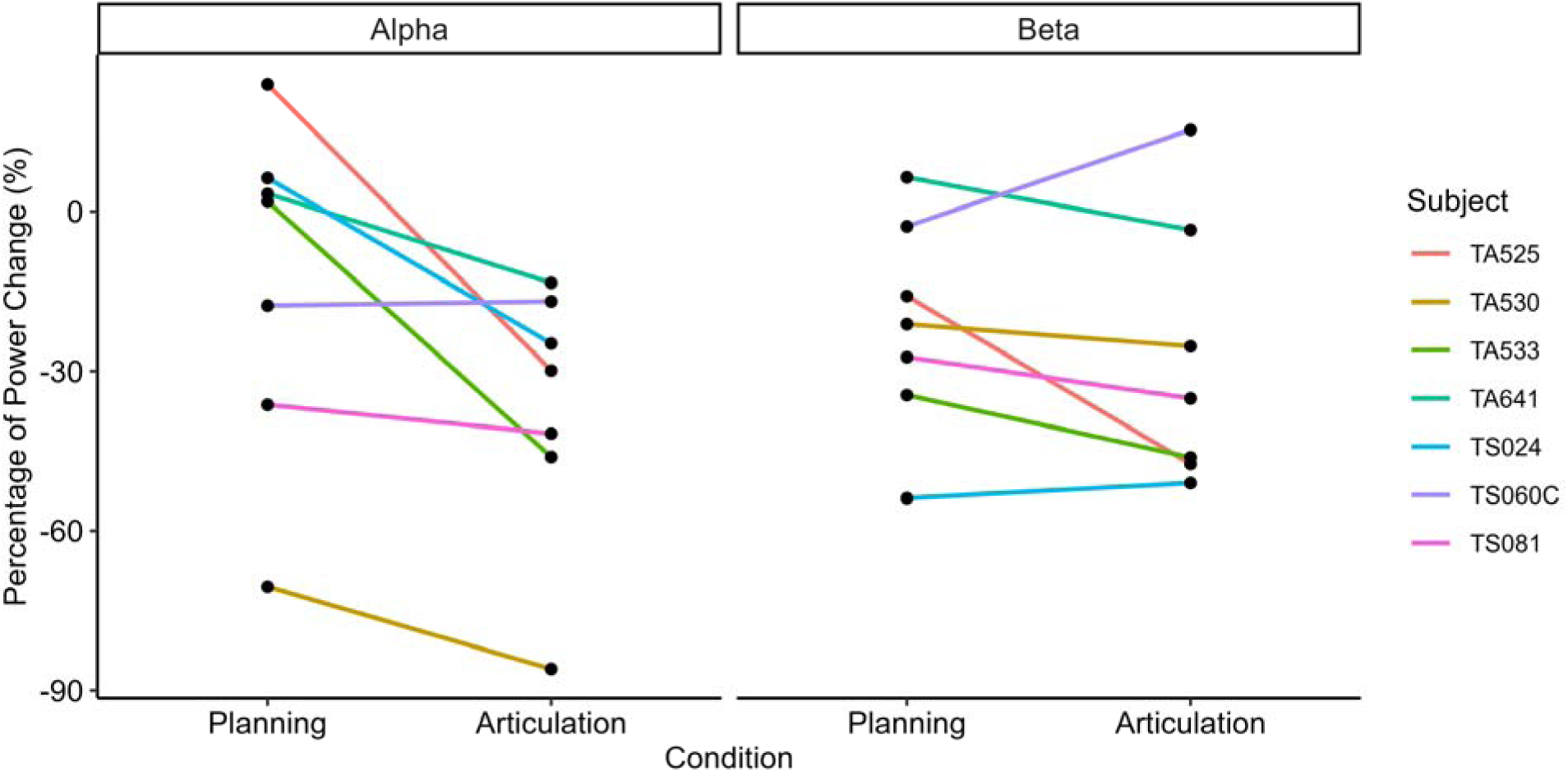
Somatosensory Alpha and Beta Power Change. Alpha (left) and beta (right) power changes in percentage relative to baseline in the speech planning and articulation stages. Each line indicates one patient.

Alpha power did not decrease significantly relative to baseline during the planning stage with anecdotal evidence (BF_10_ = .54, *r* < .01), but it did decrease during the articulation stage with strong evidence ((BF_10_ = 16.72, *r* = 1.25). Regarding our specific hypothesis, alpha power decreases (relative to baseline) were stronger (i.e., power was further decreased) during articulation than during speech planning (planning: mean = −12.72%, SD = 31.86, articulation: mean = −36.97%, SD = 24.71). Our a-priori hypothesis was supported: The data were 7.32 times more likely under the directional alternative hypothesis than under the null, suggesting moderate evidence in favor of stronger alpha ERD during articulation than during planning, with a median effect size of .91.

Beta power decreased relative to the baseline throughout the planning stage with moderate evidence (BF_10_ = 3.15, *r* = .80), and throughout the articulation stage with moderate evidence (BF_10_ = 6.16, *r* = .84). Regarding our specific hypothesis, beta ERD in the planning stage remained at a similar level as during articulation, with a slight numerical decrease on average (planning: mean = −21.31%, SD = 20.05; articulation: mean = −27.60%, SD = 25.09). The data were 1.76 times more likely under the null hypothesis (i.e., our a-priori hypothesis of no differences between planning and articulation) than under the alternative hypothesis of differential ERD between planning and articulation stages, with a median effect size of 0.31. This level of evidence, however, is considered anecdotal (Goss-Sampson et al., 2020).

### 2.3 Discussion

Using existing iEEG data to test hypotheses derived from our systematic review, we aimed to study somatosensory alpha and beta oscillations related to speech production. Consistent with our hypothesis, we found moderate evidence for a significantly larger ERD during articulation compared to the planning stage. Alpha ERD’s sensitivity to speech movements in S1 suggests its relation to somatosensory feedback processing. By contrast, only anecdotal evidence was found that beta ERD remained relatively stable during the planning and articulation stages in S1. Despite this effect being in the predicted direction, the weak nature of this evidence precludes us from firmly interpreting the role of beta ERDs in S1.

Our findings of decreased power in alpha and beta oscillations are consistent with previous extracranial and intracranial studies (e.g., Bowers et al., 2019; Bullock et al., 2023). Notably, our findings provide intracranial evidence for the association of alpha ERD with somatosensory feedback processing during speech articulation, in line with similar findings for hand movements (Stolk et al., 2019). Importantly, not only did we address the methodological concerns regarding the analysis of oscillations (Donoghue et al., 2020), we also validated the existence of ERDs in both the speech planning and speech articulation stages and found differential findings compared to the extracranial studies.

Two limitations of this study need to be mentioned here. First, the sample size was small due to the scarcity of available contacts in the ROI. That said, previous intracranial studies with the same ROI utilized samples as small as three patients (Bouchard et al., 2013), and the findings have been replicated by follow-up studies (Chartier et al., 2018; Conant et al., 2018). Second, due to the intrinsic property of open datasets, we did not have access to speech recordings and, with that, articulation offset measurements. The lack of access to articulation offset data may have introduced a confound. It is widely observed that alpha and beta power in our ROI quickly return to baseline after articulation offset (e.g., Bowers et al., 2019; Kittilstved et al., 2018), with a rebound of beta power (De Nil et al., 2021). These power increases could potentially inflate our estimation of power in the articulation stage. To minimize this potential confound effect, we only used trials with bi-syllabic word production (excluding monosyllabic word production) and we applied a conservative time window of 400 ms long, which has shown to be a reasonable duration for most bi-syllabic word production (Jacewicz et al., 2009, 2010). To reach a firmer conclusion regarding our proposed hypotheses, future studies would benefit from comprehensive contact coverage and more detailed behavioral data.

## 3 General Discussion and Conclusion

To summarize, our systematic review of existing electrophysiological studies on speech production highlight alpha and beta power decreases, or ERDs, in sensorimotor areas related to speech motor processes. These ERDs are sensitive to different functional demands, namely somatosensory feedback and motor processes. The review results led to two hypotheses: 1) function-wise, sensorimotor alpha ERD is associated with somatosensory feedback processing, whereas the beta ERD is related to motor processes; 2) considering the canonical roles of the brain areas involved, these functional modulations of alpha and beta oscillations should occur at S1 and M1, respectively. We then tested the part of the hypothesis concerning S1 using iEEG picture naming data. Our results showed (moderate) evidence for alpha oscillation modulation related to somatosensory feedback processing and (anecdotal) evidence for beta involvement in S1, supporting our hypothesis.

This study has a number of strengths. Given the excellent temporal resolution of the electrophysiological signal, we were able to clearly dissociate the speech planning from the speech articulation stages for both the systematic review and iEEG investigation. The systematic review of the literature provided not only a summary of the current literature but also provided a solid background on which to devise the hypotheses to test. We then tested part of our hypotheses using data with high spatio-temporal resolution and signal-to-noise ratio. More importantly, our analysis was deliberately controlled and focused on the rhythmic part of neural oscillations by testing the existence of the oscillations, validating their frequency boundaries, and disentangling the aperiodic influence via FOOOF (Donoghue et al., 2022). Despite the lack of M1 data in our own study, a recent iEEG study has found supporting evidence for our hypothesis. Noteworthily, Figure 3C in Bullock et al. (2023) shows strong beta ERD earlier than alpha ERD in M1 (or PreCG, authors’ own terminology).

Besides the ventral SMC area highlighted in this study, the dorsal SMC area has been shown to be involved during speech movements in various studies (e.g., Jarret et al., 2022; Kingyon et al., 2015). This area is hypothesized to support laryngeal processing based on the homunculus distribution of S1 (Belyk & Brown, 2017; Jarvis, 2019). Considering that this area is highly functionally specialized, we expect that our hypothesis may also apply to this area.

In conclusion, our systematic review suggests that alpha and beta power decreases around SMC in speech motor processes may be functionally and anatomically dissociable, with alpha ERD associated with somatosensory feedback processing in S1 and beta ERD being related to motor processes in M1. Our iEEG study tested the functional hypothesis of alpha and beta ERD in S1. The results confirmed the hypothesis derived from the systematic review that alpha ERD in S1 is responsive to somatosensory feedback processing. The hypothesis that beta ERD, associated with motor processes, should remain stable during planning and articulation in S1 was tentatively confirmed. Future studies with larger electrode coverage are needed to properly test the full hypothesis derived from the systematic review.

## Acknowledgements

Data used to perform this analysis were collected with support from the National Institutes of Health under award numbers DC014589 and NS098981 and were accessed from the Data Archive for the BRAIN Initiative with support from the National Institutes of Health under Award Number R24MH114796.

## CRediT

Lydia Z. Huang: Conceptualization, formal analysis, visualization, writing – original draft, writing – review and editing

Yang Cao: Methodology, supervision

Esther Janse: Methodology, writing – review and editing

Vitória Piai: Conceptualization, methodology, supervision, validation, writing – review and editing

## Supplementary material

Note. The bold search term in Table S1 was selected for the detailed search in Table S2. The bold numbers in Table S2 represent the studies that entered our screening procedure.

**Table S1.** Initial Search for TFS Studies.

| Search Terms | Type | Categories | Web of Science | PubMed | Google Scholar |
| --- | --- | --- | --- | --- | --- |
| motor AND “speech production” | All fields (topic) | Neuroscience (693); clinical neurology (186); neuroimaging (106); | 1754 (1678) | 2495 | 115000 |
| ~ AND neuroimaging AND “speech production” | All fields (topic) |  | 156 (111) | 489 | 22700 |
| ~ AND “oscillat***” AND “speech production” | All fields |  | 53 | 55 | 9950 |
| ~ AND electrophysiology AND “speech production” | All fields |  | 2 | 13 | 11000 |
| ~ AND “brain rhythm*” AND “speech production” | All fields |  | 1 | 47 | 21700 |
| ~ AND EEG AND “speech production” | All fields |  | 65 | 132 | 13600 |
| ~ AND MEG AND “speech production” | All fields |  | 31 | 26 | 7460 |
| ~ AND iEEG AND “speech production” | All fields |  | 1 | 1 | 177 |
| ~ AND ECoG AND “speech production” | All fields |  | 20 | 39 | 1240 |

**Table S2.** Detailed Search for TFS Studies.

| Search Terms | Type | Web of Science | PubMed |
| --- | --- | --- | --- |
| motor AND “oscillat***” AND “speech production” | All fields | **53** | **55** |
| ~ AND “language production” | All fields | **13** | **43** |
| ~ AND articulation | All fields | **25** | 122 |
| ~ AND “neural oscillation” AND articulation | All fields | 2 | **30** |
| ~ AND “vowel production” | All fields | **2** | **12** |
| ~ AND “consonant production” | All fields | 0 | **8** |
| ~ AND “syllable production” | All fields | **3** | **17** |
| ~ AND “word production” | All fields | **14** | **11** |
| ~ AND “sentence production” | All fields | 0 | **4** |
|  | Combined all above searches | **85** | **89** |

**Table S3.** Search for PWS-TFS Studies.

| **Search Terms** | **Type** | **Web of Science** | **PubMed** | **Google Scholar** |
| --- | --- | --- | --- | --- |
| neuroimaging AND stuttering | All | 96 (83) | 179 | 8820 |
|  | fields<br>(topic) |  |  |  |
| **“oscillat\*\*\*” AND ~** | **All**<br>**fields** | **46** | **31** | 9450 |
| electrophysiology AND ~ | All<br>fields | 9 | 16 | 5550 |
| “brain rhythm*” AND ~ | All<br>fields | 1 | 32 | 23000 |
| EEG AND ~ | All<br>fields | 57 | 154 | 10200 |
| MEG AND ~ | All<br>fields | 20 | 11 | 3850 |

Note. iEEG/ECOG/SEEG and “stuttering” returned no results in the first two databases, but 46, 305, and 73 for google scholar.

**Figure S1.**
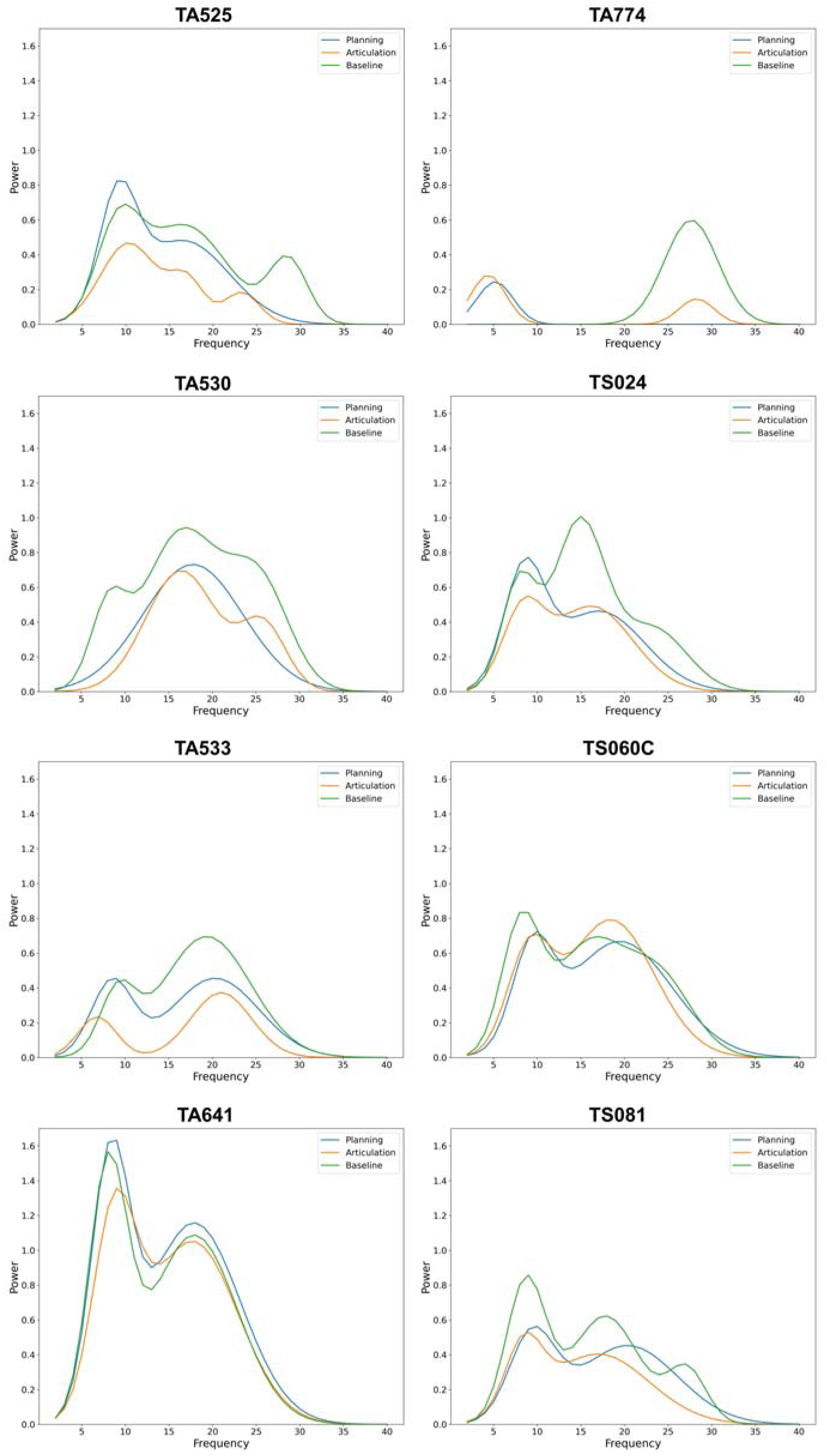
Individual Power Spectra. Fitted power spectra from FOOOF for each participant.

